# Tuberculosis in farm workers exposed to dairy and beef livestock in Colombia

**DOI:** 10.1101/821751

**Authors:** Catalina Muñoz, Johana Rueda, Luz E. Botero, Gloria I. Mejía, Ximena Cardona, Manuel G. Jaramillo, Jaime Robledo

## Abstract

The objective of this study was to determine the frequency of tuberculosis (TB) in workers from dairy and beef livestock farms in the northern part of Colombia. Tuberculin skin test and an interferon-gamma release assay (IGRA) were used for diagnosis of latent tuberculosis; sputum samples were taken from respiratory symptomatic subjects, microbiological and molecular tests were done for diagnosis of active TB. Absolute frequencies, percentages, and crude prevalence ratios were calculated, and a robust Poisson Model with adjustment by municipality was made. In 674 farm workers, latent TB frequency was 35.8%. Variables such as having had pulmonary TB (PR 2.82, 95% CI 1.90 – 4.17), having been in contact with people with active TB (PR 1.57, 95% CI 1.24 – 1.98), and having performed some undergraduate or postgraduate study (PR 1.6, 95% CI 1.03 – 2.49), were significantly associated with latent TB. No active tuberculosis disease was confirmed in symptomatic respiratory patients. The exposure level to cattle was not significantly associated with latent TB infection. In conclusion, in the studied population exposure to cattle was not a risk factor for TB, other factors commonly found in general population exposed to human TB were demonstrated.

**Author summary:** Zoonotic TB is a disease caused by the transmission of the *M. bovis* bacteria that is part of the Mycobacterium tuberculosis Complex, through contact with cattle to humans, by the consumption of unpasteurized dairy products from infected animals or by inhalation of aerosols exhaled by sick animals.

This study investigated the frequency of TB in human population related to cattle, in order to determine if there were risk factors related to TB infection or disease. Finding that there was no significant relationship between being exposed to cattle and having latent TB. However, the results of this study together with other research reported in the literature suggest that research on zoonotic and bovine TB should be continued, especially about epidemiology, diagnostic methods, health systems and interventions coordinated with veterinary services.

## Introduction

Tuberculosis (TB) is the ninth cause of death and the first cause of death due to infectious diseases worldwide. In 2017, ten million people became ill with TB and 1.3 million died from this disease [1].

Approximately one-quarter of the world’s population (1.7 billion) has latent TB, that is, they are infected with the bacillus but have not yet become ill or can transmit the infection; 5 to 15% of this population will develop an active disease throughout life [2]. The National Public Health Surveillance System of Colombia (SIVIGILA) reported 14,420 new cases of TB in 2018, out of which 2,609 cases were said to have occurred in Antioquia [3].

Tuberculosis is caused by diverse species belonging to *Mycobacterium tuberculosis* complex (CMTB). In humans, the species that cause most cases is *Mycobacterium tuberculosis* (MTB), followed by *Mycobacterium bovis,* which is, in turn, the causative agent of bovine TB in animals and of zoonotic TB in humans [1,4]. Cattle are the definitive hosts for *M. bovis,* but other domestic and wild mammals can also be infected. The transmission of bovine TB to humans is mainly due to the consumption of unpasteurized dairy products from infected animals or by the inhalation of aerosols exhaled by diseased animals. Less common forms to transmit the bovine TB are the consumption of undercooked contaminated beef and direct percutaneous contact, associated mainly with the bacteria infecting a wound [5].

Currently, the importance of studying zoonotic TB in the context of *One Health* approach has facilitated the implementation of health measures to improve food safety, such as the general pasteurization of milk and the control of bovine TB in animal reservoirs that improves its control [6]. However, the risk of disease in humans persists particularly in specific occupational categories in the livestock sector that require close and constant contact with potentially infected animals as zootechnicians, milkers, livestock managers, and staff at animal management plants [7].

According to the Colombian Agricultural Institute (ICA), there are confirmed cases of *M. bovis* in bovines every year; this means a potential risk of infection with tuberculosis for people associated with these animals. However, the real situation of *M. bovis* in livestock in the country is mostly unknown. In Colombia, epidemiological surveillance of human TB is restricted to the identification of CMTB but not to the differentiation of the species that cause the disease, which does not allow to identify human cases of TB caused by *M. bovis*. All of the above makes it necessary to know better the contribution of bovine TB to human TB cases in the country that allow re-evaluating current health policies, generate better health controls, and educate the population in this economic sector. Antioquia is a department located in the northwest of Colombia. Part of its economy has developed around livestock activity, and it currently has the highest livestock production in the country, with 11.75% of the national bovine population [8]. So far, there is no information on the frequency of TB in people exposed to cattle in Antioquia.

This study aimed to determine the frequency of TB in a human population exposed to dairy and beef livestock in Antioquia, Colombia.

## Methodology

An analytical cross-sectional study was carried out between February 2017 and February 2018. The study involved people who worked on farms associated to *Colanta*® a cooperative of dairy farms and the “National program for the control and eradication of bovine tuberculosis and the certification of farms free of this disease” offered by the ICA, in addition, technical staff from other institutions in the agricultural area located in Antioquia was included.

### Characteristics of the population

The study included participants over 18 years old, who had direct or indirect contact with dairy and beef livestock. Individuals who were unable to attend the tuberculin skin test (TST) read after 48 to 72 hours, who did not respond adequately to the clinical-epidemiological survey and those who refused to participate in the study were excluded (Fig. 1).

**Figure 1.**
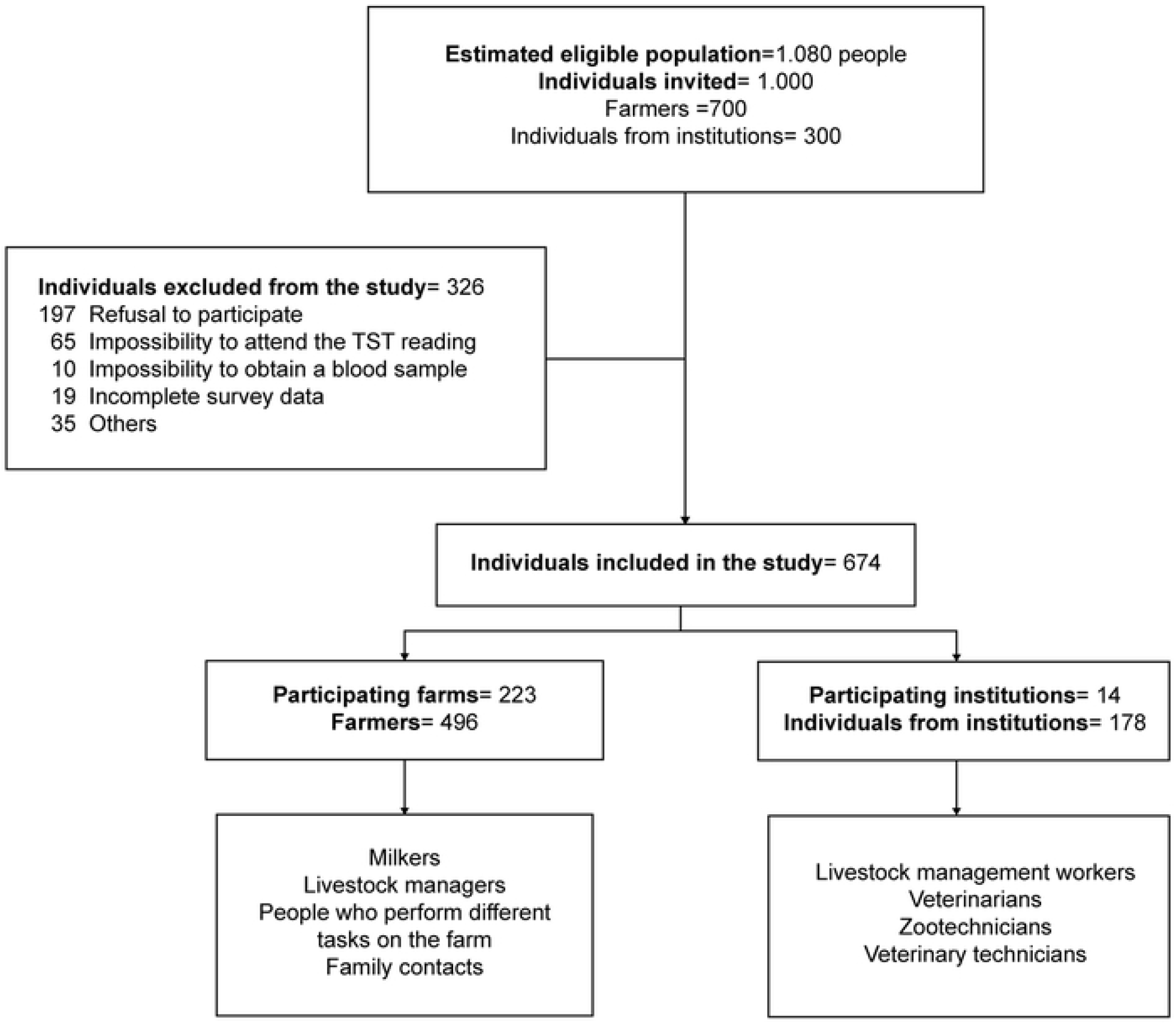
Admitted and analyzed population

All participants completed a clinical-epidemiological survey and signed two informed consents: one regarding the sampling of venous blood for the detection of interferon-gamma (IFN-γ) with QuantiFERON®-TB GOLD PLUS (QFT^®^-Plus), and the other, regarding the application of the TST test.

### Exposure level

The exposure level of the population to cattle were stratified according to a published work [9] and adjusted accordingly, as follows: (i) High degree of exposure (Individuals directly exposed to cattle, in full-time works, mainly indoors.) (ii) Average degree of exposure: (Individuals directly exposed to cattle, in part-time works not exclusive to livestock, usually outdoors. (iii) Low degree of exposure: Individuals occasionally and indirectly exposed, during one hour or less, that usually perform tasks unrelated to cattle handling).

### Latent tuberculosis diagnosis

A QFT^®^-Plus test [10] was performed. The amount of IFN-γ (IU/ml) present in the measuring tubes due to the reaction of ESAT-6 and CFP-10 proteins (antigens associated with CMTB) was measured with an ELISA method [10]. Also, the TST test was performed using the Mantoux method [11], with an intradermal injection of 5 IU (0.1 mL) of purified protein derivative (Manufactured by BB-NCIPD-LTD 1504 Sofia, Bulgaria) on the forearm. After 72 hours, the reading was conducted by previously trained personal. The results were recorded in millimeters and considered positive when the induration was equal to or greater than 10 millimeters [12,13]. According to WHO, a positive result on TST or IGRA leads to a diagnosis of latent TB [14].

### Active tuberculosis diagnosis

A sample of sputum was collected from symptomatic respiratory patients (SR) [15], Decontamination and liquefaction processes were carried out per sample, using the N-acetyl-L-cysteine method, together with 4% sodium hydroxide [16]. The sample was evaluated microscopically with bacilloscopy with Auramine-Rhodamine stain and then inoculated in solid (Lowenstein Jensen) and liquid (MGIT Becton-Dickinson tubes, Bactec MGIT 960^®^, Sparks, MA, USA) culture medium for microorganism recovery and subsequent identification. A molecular test was also conducted with all sputum samples using the Xpert MTB/Rif^®^ method to detect the CMTB bacteria. Participants were also evaluated to identify other clinical manifestations that may indicate extrapulmonary TB.

### Statistical analysis

The information obtained from the clinical-epidemiological survey was tabulated in a Microsoft Excel 2016 database. This information was transferred to the IBM SPSS Statistics 25.0 and Stata program 15.0 to perform the necessary statistical analyses.

Absolute frequencies and percentages were used for the descriptive analysis of qualitative variables. To evaluate the variables that were associated with the latent TB positivity, crude prevalence ratios were used, and a robust Poisson Model was made with a standard error adjustment by municipality grouping. Variables with p<0.25 values (Hosmer and Lemeshow criteria) were included in the model. A 0.05 alpha was considered for statistically significant associations.

Agreement between QFT^®^-Plus and TST tests was assessed by using Kappa statistic (k). It was considered that a ‘poor’ agreement was κ ≤ 0.20, an ‘acceptable’ agreement was 0.20 <κ ≤ 0.40, a ‘moderate’ agreement was 0.40 <κ ≤ 0.60, a ‘good’ agreement was 0.60 <κ ≤ 0.80, and a ‘very good’ agreement was 0.80 < κ ≤ 1. [17].

### Ethical considerations

According to Colombian Resolution No. 8430 of 1993, the individuals participating in this study were exposed to minimal risk. The Research Ethics Committee of the Corporation for Biological Research and the Research Ethics Committee of the School of Health Sciences of the Universidad Pontificia Bolivariana in Medellín, Colombia approved the study. This study had research purposes and sought the collective benefit of the community directly involved in livestock activities of the scientific community and the general population.

## Results

The eligible population was constituted by 1080 individuals located in the department of Antioquia. Out of these, 1000 were invited to participate. Seven hundred people (700) (74%) came from different farms in the department, and 300 (26%) came from institutions responsible for the production, monitoring, and control of dairy and beef production. The 326 individuals excluded did not meet the inclusion criteria or decided not to participate. In total, data from 674 individuals were analyzed (Figure 1).

The age of the participants ranged from 18 to 83 years, 79.6% were men, and 44.2% of the participants in the study went to primary school (Table 1).

**Table 1.** Demographic characteristics of the population (N=, 674).

| VARIABLE | n | % |
| --- | --- | --- |
| Males | 537 | 79.6 |
| Age |  |  |
| ≤34 | 208 | 30.9 |
| 35 - 49 | 265 | 39.2 |
| 50 - 64 | 163 | 24.2 |
| ≥65 | 38 | 5.6 |
| Educational Level |  |  |
| None | 32 | 4.7 |
| Primary School | 298 | 44.2 |
| High School | 183 | 27.2 |
| Technical or Technological education | 59 | 8.8 |
| Undergraduate or Postgraduate | 102 | 15.1 |

N: population size, n: sample size

Regarding the geographical distribution of the population, 51.6% came from the northern region of the Department of Antioquia (Fig. 2), specifically the municipalities of San Pedro de los Milagros, Entrerríos, Santa Rosa de Osos, Don Matías, Yarumal, Belmira and San José de la Montaña.

**Figure 2.**
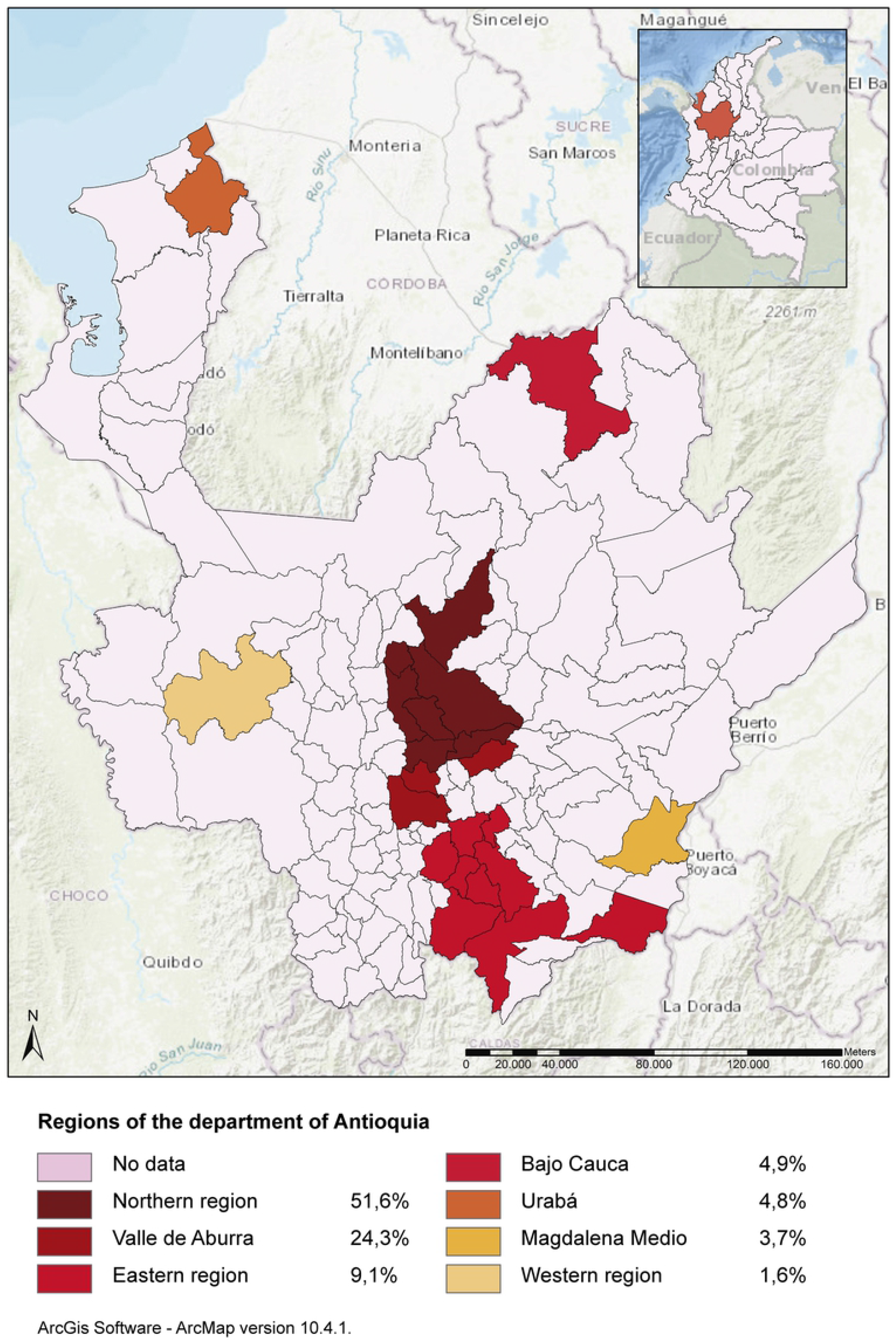
Distribution of the population of study according to their regions of origin in Antioquia, Colombia.

Most participants, 86.8% had been vaccinated with BCG, 6.8% reported previous close contact with a case of pulmonary TB, and 0.9% reported to have been previously diagnosed with TB. Most of the population, 85.6%, were directly exposed to cattle, and 69% reported being exposed to sick cattle. Out of all participants, 41.2% reported consumption and preparation of unpasteurized dairy products, and 95.8% consumed beef (Table 2).

**Table 2.** Clinical-epidemiological characteristics of the population (N=674).

| VARIABLE | n | % |
| --- | --- | --- |
| BCG vaccine | 585 | 86.8 |
| Previously diagnosed TB | 6 | 0.9 |
| Contact with people with TB | 46 | 6.8 |
| Time spent working on a farm or institution (months) |  |  |
| ≤24 | 176 | 26.1 |
| 25 - 96 | 162 | 24.0 |
| 97 - 240 | 189 | 28.0 |
| >241 | 147 | 21.8 |
| Direct exposure to cattle | 577 | 85.6 |
| Exposure to sick cattle* | 465 | 69.0 |
| Use of mask during cattle care | 119 | 17.7 |
| Exposure level to cattle |  |  |
| Low | 41 | 6.1 |
| Medium | 106 | 15.7 |
| High | 527 | 78.2 |
| Raw milk consumption | 206 | 30.6 |
| Products derived from raw milk |  |  |
| No consumption – No preparation | 291 | 43.2 |
| Consumption or preparation | 105 | 15.6 |
| Consumption and preparation | 278 | 41.2 |
| Beef consumption | 646 | 95.8 |
| Beef cooking term (N=646) |  |  |
| Undercooked beef | 72 | 11.1 |
| Cooked | 128 | 19.8 |
| Well cooked | 446 | 69.0 |

N: Population size, n: Sample size
\*Sick cattle with any type of condition.

Positive results by any of the latent TB tests performed (TST and QFT^®^-Plus) were in 35.8% of individuals, 32.9% tested positive with TST and 10.7% with QFT^®^-Plus (Table 3). Out of the total population, 8% tested positive in both tests, (Kappa coefficient was 0.237).

**Table 3.**
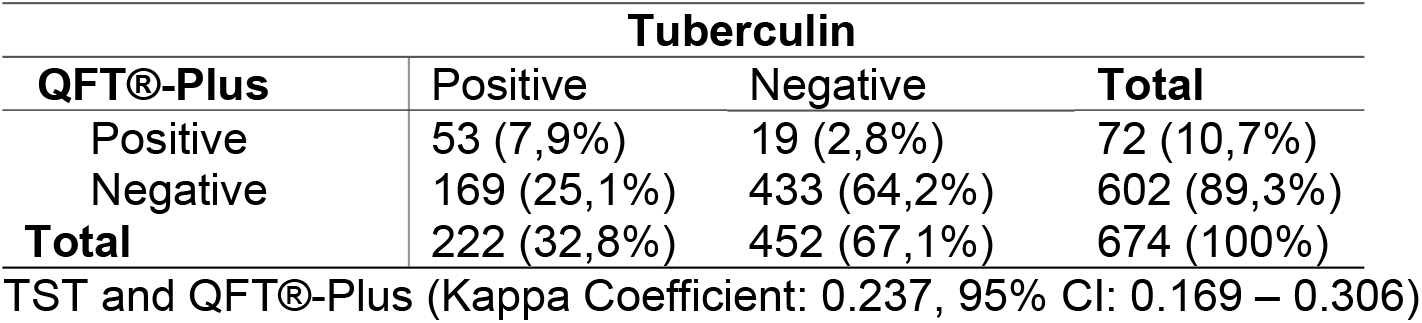
Agreement between TST and QFT^®^-Plus (N=674)

Significantly associated variables with latent TB were: prior diagnosis of TB (PR 2.82, CI 95% 1.90 – 4.17) contact with people with TB (PR 1.57, CI 95% 1.24 – 1.98), and completion of undergraduate or postgraduate studies (PR 1.6, CI 95% 1.03 – 6 2.49). (Table 4).

**Table 4.** Association of sociodemographic and epidemiological variables to latent tuberculosis in the study population (N=674).

| VARIABLES | n/N (%) | PR<br>(raw) | CI 95% |  | PR*<br>(adjusted) | CI 95% |  | Value<br>*p* |
| --- | --- | --- | --- | --- | --- | --- | --- | --- |
| Sex |  |  |  |  |  |  |  |  |
| Female | 43/138 (31.2) | 1 |  |  |  |  |  |  |
| Male | 197/536 (36.8) | 1.18 | 0.85 | 1.63 | 1.32 | 0.97 | 1.77 | 0.062 |
| Age (in groups) |  |  |  |  |  |  |  |  |
| <34 | 63/208 (30.3) | 1 |  |  | 1 |  |  |  |
| 35 - 49 | 106/265 (40.0) | 1.32 | 0.98 | 1.77 | 1.34 | 0.99 | 1.81 | 0.052 |
| 50 - 64 | 60/163 (36.8) | 1.22 | 0.86 | 1.72 | 1.25 | 0.91 | 1.72 | 0.171 |
| ≥65 | 11/38 (28.9) | 0.96 | 0.57 | 1.60 | 1.03 | 0.66 | 1.60 | 0.897 |
| Education level |  |  |  |  |  |  |  |  |
| None | 8/32 (25.0) | 1 |  |  | 1 |  |  |  |
| Primary school | 97/298 (32.6) | 1.3 | 0.84 | 2.02 | 1.11 | 0.73 | 1.69 | 0.617 |
| High school | 70/183 (38.3) | 1.53 | 0.90 | 2.60 | 1.4 | 0.85 | 2.33 | 0.191 |
| Technical or technological education | 20/59 (33.9) | 1.36 | 0.79 | 2.34 | 1.22 | 0.75 | 1.99 | 0.420 |
| Undergraduate or Postgraduate | 45/102 (44.1) | 1.76 | 1.15 | 2.70 | 1.6 | 1.03 | 2.49 | 0.037** |
| Previously diagnosed TB |  |  |  |  |  |  |  |  |
| No | 234/668 (35.0) | 1 |  |  | 1 |  |  |  |
| Yes | 6/6 (100.0) | 2.85 | 2.40 | 3.40 | 2.82 | 1.90 | 4.17 | <0.001*<br>* |
| Contact with people with TB |  |  |  |  |  |  |  |  |
| No | 215/628 (34.2) | 1 |  |  | 1 |  |  |  |
| Yes | 25/46 (54.3) | 1.59 | 1.31 | 1.92 | 1.57 | 1.24 | 1.98 | <0.001*<br>* |
| Direct exposure to cattle |  |  |  |  |  |  |  |  |
| No | 31/97 (32.0) | 1 |  |  | 1 |  |  |  |
| Yes | 209/577 (36.2) | 1.13 | 0.68 | 1.90 | 1.64 | 0.89 | 3.02 | 0.111 |
| Raw milk consumption |  |  |  |  |  |  |  |  |
| No | 162/468 (34.6) | 1 |  |  | 1 |  |  |  |
| Yes | 78/206 (37.9) | 1.09 | 0.88 | 1.36 | 1.15 | 0.91 | 1.45 | 0.240 |

PR: Prevalence Ratios, N: Population size, n: sample size, CI: Confidence Interval
\*Robust Poisson model with error adjustment by municipality
\*\* $p < 0.05$

The variable “Exposure Level” was classified as a variable of occupational exposure to cattle. There was no statistically significant association with the variable “Latent TB” (value p-0.291) (Table 5).

**Table 5.** Association between the exposure level to cattle and the positivity of tests for latent TB

| VARIABLE |  | Positive Results - TST or QFT |  |  | Value p |
| --- | --- | --- | --- | --- | --- |
|  |  | n/N | % | IC 95% |  |
| Exposure level to cattle | **Medium** | 32/105 | 30.5 | 1 | 0.291 |
|  | **High** | 191/528 | 36.2 | 1.44 (0.92 – 2.26) |  |
|  | **Low** | 18/41 | 43.9 | 1.18 (0.87 – 1.61) |  |

N: Population size, n: sample size, CI: Confidence Interval

Twenty-four (3.6%) participants included in the study had respiratory symptoms compatible with TB. The studies did not find positivity for CMTB microorganisms in the microbiological and molecular tests of SR’s sputum samples; therefore, not active pulmonary TB cases were confirmed. Tests related to the symptomatology of individuals, aimed at detecting pathogens other than mycobacteria were not performed.

## Discussion

Human TB is still a problem worldwide, and Colombia is not an exception. Most cases are caused by *M. tuberculosis*, but *M. bovis* has been identified as a cause of TB in humans in different regions of the world [2]. According to the latest WHO report, approximately one-quarter (25%) of the world’s population has latent TB (2). The present study found that approximately one-third (35.8%) of the population exposed to bovines is infected; However, the prevalence of latent TB in other Colombian populations is higher. A study conducted by Ochoa and collaborators in 2017 [18] showed that, in Medellin, the prevalence of latent TB in health staff was 62%. Likewise, a study conducted by del Corral and collaborators in 2009 [19] found that the prevalence of latent TB was 65,9% among household contacs of TB patients compared with 42.7% of latent TB of the general population in a population in urban location in Medellín.

In this study, 674 people were analyzed. Out of these, 32.9% tested positive with TST test and 10.7% with IGRA. Compared to other studies in similar populations, the frequency of latent TB was lower. Torres-González and collaborators found that in a group of 311 dairy farmworkers in Mexico, the prevalence of latent TB was 76.2% with TST and 58.5% with IGRA [9]. Likewise, Oren and collaborators analyzed 109 agricultural migrant workers from the US-Mexico border. They found that the prevalence of latent TB was 34% with TST and 50% with IGRA [17]. One of the possible reasons for the low frequency of latent TB found in this study may be the selected population. The farms under certification for bovine tuberculosis were 91%, that is to say, being free of bovine TB according to the national policies. In this context, the human population studied would have a lower risk of becoming infected. On the other hand, a significant association was found between a previous contact of people with active pulmonary TB patients and having latent TB, which confirm data from studies in general population [20,21].

Only 8% of the study population tested positive with TST and QFT^®^-Plus simultaneously, demonstrating a ‘regular’ agreement (k-0.24) between the two tests. These findings are consistent with other studies. In 2013, a study by Jo and collaborators [22] found that the agreement between these tests was ‘regular’ (k-0.22); in 2017, Ochoa and collaborators [18] found a ‘moderate’ agreement (k-0.47) between these two tests on health care workers. According to WHO guidelines, regardless of the concordance between TST and QFT, both tests could be used to diagnose latent TB; however, the findings of this study do not allow us to conclude which of the two tests is more advisable to make a diagnosis of latent TB in people exposed to cattle.

The frequency in males was 79.6% (537/674), given that most people who carry out activities associated with livestock production are usually male. This is coherent with other studies where the frequency of males was also high [9,17,23–25]. Likewise, according to WHO, worldwide, men have a significantly higher risk of contracting TB compared to women [26]. However, when analyzing the relationship between gender and the presence of latent TB, no significant association was demonstrated.

In relation to the level of education, the primary school was the highest educational level that a large proportion of individuals had reached (44.2%). This is common in rural contexts, where people usually start working from a very early age to support their families and increase incomes. However, 15% of participants carried out undergraduate and graduate studies. A statistically significant association between this academic level and latent TB was found, which may be explained by the fact that professionals included in the study were mainly veterinary doctors and zootechnicians. These individuals are constantly and directly exposed to cattle, which may be indicative of greater exposure to sick animals and a higher possibility of infection with mycobacteria.

According to the time spent working with cattle, 28% (189/674) of the participating population had been carrying out livestock activities for more than 8 years (97 and 240 months). Although this should increase the likelihood of becoming infected with CMTB bacteria, no significant association between time working with cattle and having latent TB was demonstrated. Nevertheless, data obtained by Torres and collaborators indicated that habits of working with animals constitute a significant risk of having latent TB [9].

Previous studies have shown that the main source of transmission of zoonotic TB is the consumption of unpasteurized milk and its derivatives [27]; however, this study did not show that there is a risk of having latent TB with the consumption of unpasteurized products. Even so, the importance to continue educating people to eat appropriately pasteurized foods should be stressed. It is worth considering that some of the participants may have deliberately denied the consumption of raw milk given that farm owners usually prohibit this type of activity because of the risk that milk-borne pathogens represent [28].

A low percentage of the population involved (17.7%) reported the use of a mouth mask when working with cattle. Despite this being an essential protective measure to avoid the transmission of the microorganism, no statistically significant association with latent TB was found [5].

The risk of latent TB was also found in individuals who reported to have been treated for active TB before the study (0.9% - 6 individuals out of 674). TST and IGRA tests remained positive in these cases because of the positive hypersensitivity response evidenced through TST after exposure to the agent [29] and the fact that IGRA can also remain positive for a considerable period in people who have received treatment for the disease [30].

Twenty-four (3.6%) of the 674 participants were listed as SR. Microbiological and molecular analyses did not demonstrate the presence of *M. bovis* in any individual, even though it is the bacteria expected to circulate in this type of population. Nevertheless, the findings revealed a higher prevalence of latent TB in the population of study than in the general population. The tests used to diagnose latent TB did not allow us to determine whether the infection is caused by *M. bovis* but demonstrated that there is a transmission of CMTB bacteria in this population. As it cannot be concluded that latent TB is transmitted by cattle, there is a need to seek and develop new techniques aimed at detecting latent TB caused by *M. bovis*.

The limitations of this study include the impossibility to perform the second TST test to detect the booster effect recommended by WHO, which increases TST positivity between 10 and 15% [31,32]. Most of the selected participants belonged to farms involved in the TB program promoted by the ICA, whose characteristics impede the extrapolation of results to the general population exposed to livestock. For this reason, it is necessary future research that involves individuals working with livestock from the general population, in order to see if the characteristics of the population vary or remain and to understand the impact of exposure to cattle on human health, particularly in regards to TB.

In conclusion, this study did not find a significant relationship between exposure to cattle and active or latent TB. Our results suggest that people who adhere to bovine TB programs may have better protective measures when working with cattle. Further research in the field of zoonotic and bovine TB is required, especially concerning its epidemiology. It is also essential to study other animals in wildlife reservoirs that may expand the scope of transmission. There is also a need for new diagnostic methods and coordinated interventions with veterinary services and health systems in rural areas where public policies fail to meet their objectives.

## Acknowledgments

To the Colombian Agricultural Institute (ICA) and the Directorate of Risk Factors of the Health and Social Protection Section of Antioquia for the information and support provided, which allowed the completion of this project.

## Financing

The Administrative Department of Science, Technology, and Innovation - Colciencias, Colombia, approved the financing of the project “Prevalence of latent and active bovine tuberculosis among workers exposed to cattle on dairy farms in Antioquia, Colombia” with code 221374456133 and contract 737-2016.

